# Elucidation of ion effects on the Thermodynamics of RNA Folding

**DOI:** 10.1101/364935

**Authors:** Natalia A. Denesyuk, D. Thirumalai

## Abstract

How ions affect RNA folding thermodynamics and kinetics is an important but a vexing problem that remains unsolved. Experiments have shown that the free energy change, Δ*G*(*c*), of RNA upon folding varies with the salt concentration (*c*) as, Δ*G*(*c*) = *k_c_* ln *c* + const, where the coefficient *k_c_* is proportional to the difference in the uptake of ions (ion preferential coefficient), ΔΓ, between the folded and unfolded states. We performed simulations of a coarse-grained model, by modeling electrostatic interactions implicitly and with explicit representation of ions, to elucidate the molecular underpinnings of the relationship between folding free energy and ion preferential coefficient. Without any input from experiments, the simulations quantitatively reproduce the heat capacity for the −1 frame shifting pseudoknot (PK) from Beet Western Yellow Virus, thus validating the model. We show that ΔG(c) calculated directly from ΔΓ varies linearly with ln *c* (*c* < 0.2*M*), for a hairpin and the PK, thus demonstrating a molecular link between the two quantities for RNA molecules that undergo substantial conformational changes during folding. Explicit ion simulations also show the linear dependence of Δ*G*(*c*) on ln *c* at all *c* with *k_c_* = 2*k_B_T*, except that Δ*G*(*c*) values are shifted by about 2 kcal/mol higher than experiments at all salt concentrations. The discrepancy is due to an underestimate the Γ values for both the folded and unfolded states, while giving accurate values for ΔΓ. The predictions for the salt dependence of ΔΓ are amenable to test using single molecule pulling experiments. Our simulations, representing a significant advance in quantitatively describing ion effects in RNA, show that the framework provided here can be used to obtain accurate thermodynamics of RNA folding.

## Introduction

In common with proteins, RNA molecules that carry out cellular functions also adopt specific, compact conformations, which require ions. In the absence of counter ions, compact RNA structures are energetically unfavorable due to the close proximity of negatively charged phosphate groups. Therefore, to enable RNA molecules to fold, counterions from the buffer solution must condense onto the sugar-phosphate backbone, which would reduce the charges on the phosphate groups. The counterion condensation establishes a close relationship between RNA structures populated at equilibrium and ionic environment.^1–11^ Divalent ions, typically Mg^2+^, are particularly efficient in stabilizing RNA folded structures.^3,12–17^ Representative high resolution structures of folded RNA show individual Mg^2+^ ions are bound to multiple phosphate groups.^18,19^ But the presence of divalent ions is not essential for the stability of many RNAs with relatively simple architectures. For example, the —1 frameshifting pseudoknot from beet western yellow virus^14,20,21^ (BWYV PK, Fig. 1) is stable in relatively low concentrations of monovalent salt even in the absence of Mg^2+^. Experiments have shown that the thermodynamic stability, Δ*G*, of the BWYK PK in a Na^+^ buffer increases linearly with the logarithm of the salt molar concentration, *c*.^14^ Likewise, experimental data for the stability of RNA hairpins^22,23^ and of polymeric RNA duplexes and triplexes^24^ in monovalent salt buffers corroborate the linear dependence, Δ*G*(*c*) = –*k_c_* ln *c* + const, where the value of *k_c_* depends on the specific RNA structure. A similar relationship has been established for oligo and polymeric DNA double helices,^25,26^ for which *k_c_* is known to be largely insensitive to the DNA sequence. Furthermore, if monovalent salt buffers contain relatively low concentrations of divalent salt, *c*_2_ ≪ *c*, the observed dependence of Δ*G* on ln *c*_2_ is also found to be linear.^14^ Despite the wealth of data on these systems and theoretical and several computational studies^9,27–32^ a molecular understanding of ion effects on RNA folding is lacking. More importantly, there is practically no computational framework that could reliably predict the ion-dependent free energies and how they change as RNA folds. Here, we go a long way in solving the problem by using molecular simulations of a coarse-grained model combined with theory to predict the folding free energies of a RNA hairpin and a pseudoknot (PK) in the presence of both explicit and implicit *Na*^+^ ions.

**Figure 1:**
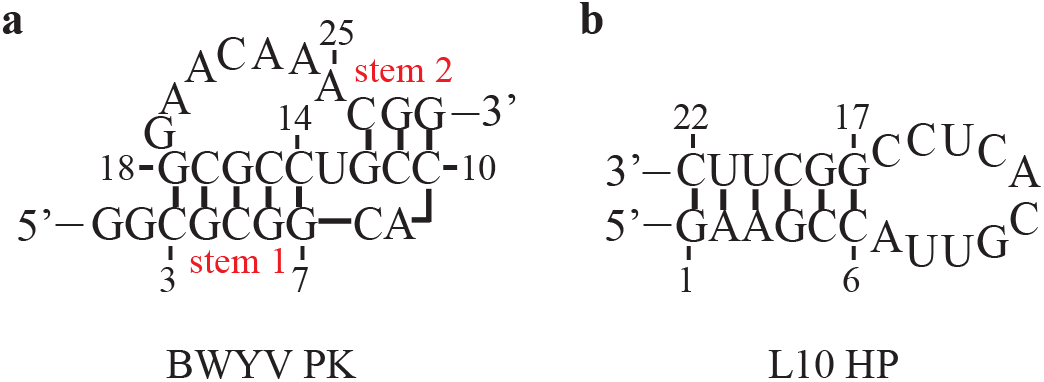
Secondary structures of studied RNA. The L10 HP contains a 5’-pG and the total of 22 phosphate groups. The BWYV PK does not have a phosphate group at the 5’-end.

Pioneering theoretical studies of the influence of ionic conditions on the stability of nucleic acid structures have linked the coefficient *k_c_* to the quantity known as the ion preferential interaction coefficient, Γ. For monovalent salts, *k_c_* = 2*k*_B_*T*ΔΓ, where ΔΓ = Γ_f_ — Γ_u_ is a difference in the preferential interaction coefficient between the folded and unfolded states of the nucleic acid molecule.^33,34^ The coefficient Γ is defined as the partial derivative,

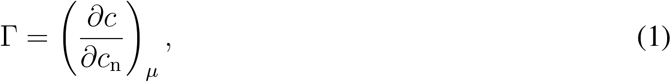

where *c*_n_ is the nucleic acid concentration and *μ* is the chemical potential of ions.^33,34^ In other words, Γ is the number of excess uptake of ions per nucleic acid molecule, as compared to the solution without nucleic acid but with the same value of *μ*. The limit *c*_n_ → 0 is assumed in Eq. (1) to avoid the influence of interactions between the polyions.

The established theoretical framework relating Δ*G*(*c*) and ΔΓ has no underlying model for the folding/unfolding transition in RNA or DNA. Instead it relies on the identity

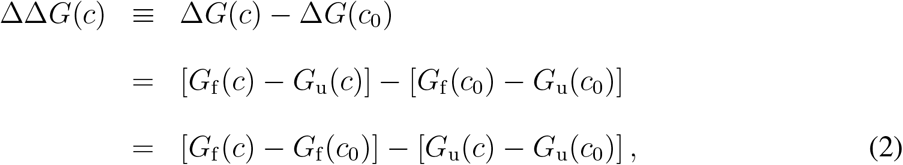

where *G*_f_ and *G*_u_ are the free energies of the folded and unfolded states, and *c*_0_ is a reference salt concentration. Using Eq. (2) one can obtain the electrostatic contribution, ΔΔ*G*(*c*), to the total stability, Δ*G*(*c*), from individual electrostatic free energies of the folded and unfolded states. No model for the transition between the two states is required. The electrostatic free energy of the folded state is given by^33,34^ 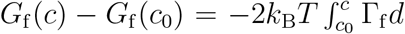 ln *c*, where *c* is the monovalent salt concentration (a similar expression holds for the unfolded state). In the case of divalent salt, the prefactor 2 on the right-hand side is omitted.^21^ It follows from Eq. (2) and the expression relating the free energy and Γ_f_ that

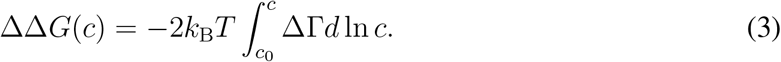

Experimental evidence that Δ*G*(*c*) changes linearly with ln *c*, implying that ΔΓ is largely insensitive to *c*.

The relationship between nucleic acid thermodynamic stability and ΔΓ has been validated for different systems using combined theoretical-experimental and purely experimental approaches. In several studies the preferential interaction coefficients of multi-stranded nucleic acid polymers were calculated from either Manning’s theory of counterion condensation^35–37^ or numerical solutions of the nonlinear Poisson-Boltzmann equation.^34^ In these theoretical treatments, duplexes, triplexes and single-stranded nucleic acids were modeled as infinitely long rigid rods characterized by three different (adjustable) linear charge densities. The resulting theoretical estimates of ΔΓ were consistent with the experimental thermodynamic data for the order-disorder transitions of polymeric DNA and RNA in monovalent salt buffers with *c* < 0.2 M. In another, purely experimental study of the BWYK PK, it proved possible to obtain accurate estimates of ΔΓ as a function of Mg^2+^ concentration using fluorescence.^21^ The curve ΔΔ*G*(*c*), which was derived from these measurements and an analogue of Eq. (3) (without the prefactor 2), agreed well with the Mg^2+^- dependent stability of the BWYK PK extracted directly from melting experiments.^14,21^

Here, we employ computer simulations to investigate the quantitative relationship between Δ*G*(*c*) and ΔΓ for small RNA molecules in monovalent salt buffers. We have previously developed a coarse-grained simulation model for RNA, which included implicit description of the ionic environment.^30^ The parameters of the model were trained by reproducing the experimentally measured melting temperatures of oligonucleotides, which in part contributes to the success of the model. We demonstrated the model is thermodynamically accurate for several RNA molecules over a wide range of monovalent salt concentration *c*, and temperature *T*. An important factor contributing to the accuracy of our modeling is its ability to capture a complete ensemble of RNA conformations, as opposed to rigid-molecule description of the folded and unfolded states. In this paper, we present the results of the same simulation model for the folding thermodynamics of the BWYV PK (Fig. 1). We obtain ΔΔ*G*(*c*) in two ways: (a) directly, from the folding/unfolding equilibrium at various c and (b) indirectly, by first computing the coefficients Γ_f_, Γ_u_ and then employing Eq. (3). We find that both approaches give the same dependence of ΔΔ*G* on ln *c*, even though the underlying dependence of ΔΓ on ln *c* is non-monotonic. In addition, we report the results of simulations of hairpin L10 (Fig. 1) using the same model as for the BWYV PK, but with explicit description of ions. In these simulations, we employ a grand canonical ensemble for ions and obtain Γ by means of direct counting of ions in the simulation box. We find that ΔΓ resulting from this counting procedure is consistent with the slope of Δ*G* vs. ln c extracted directly from our L10 melting curves. Finally, comparisons of the simulation and the observed Δ*G*(*c*) demonstrate relative advantages and limitations of implicit and explicit modeling of ions in simulations.

## Results

### Implicit ion simulations of the BWYV PK

We first discuss our simulation results for the BWYV PK (Fig. 1), whose high-resolution crystal structure is known (PDB entry 437D). The list of hydrogen bonds in the BWYV PK (Table 1) reveals an extensive tertiary structure which involves base triples. The stability of the BWYV PK tertiary structure was found to be pH sensitive and related to the protonation state of base C8.^20^ Additional experiments showed that this base was protonated in > 90% cases at pH 7.0.^14^ Assuming pH 7.0 in simulations, we added a hydrogen bond between N3 of C8 and O6 of G12 to the list of hydrogen bonds. Figure 2a shows the heat capacity of the BWYV PK obtained in experiment at 0.5 M K^+^ and pH 7.0,^20^ and in implicit ion simulations with *c* = 0.5 M and *b* = 4.4 Å. The agreement between simulation and experiment is excellent, without any adjustments to the model parameters. The model accurately predicts the two peaks in the heat capacity profile, which indicate melting of stem 1 at a higher temperature and melting of stem 2 together with the tertiary structure at a lower temperature. The finding that stem 1 melts at the higher melting temperature is in accord with the principle that assembly of PKs in general is determined by the stabilities of the constituent secondary structural elements,^38^ which in this PK are Stem 1 and Stem 2. Figure 2a also shows the predicted heat capacity of the BWYV PK at *c* = 0.05 M. The data demonstrate that the melting transition of stem 2 is more sensitive to *c* than the melting transition of stem 1. Because stem 1 is stable around the melting temperature of stem 2, the folding of stem 2 yields the native conformation of the BWYV PK. The pseudoknot is distinguished by three aligned strands of the negatively charged backbone and its stability (or melting temperature) is expected to depend strongly on the ion concentration. By contrast, the folded conformation of stem 1, the hairpin, has only two RNA strands in close contact and thus shows a weaker sensitivity to *c*.

**Table 1:** Hydrogen bonds in the BWYV PK

| Residues in contact | Hydrogen bonds |
| --- | --- |
| C3-G18 | N4-O6; N3-N1; O2-N2 |
| G4-C17 | N1-N3; N2-O2; O6-N4 |
| G4-A20 | N2-N3; O2'-N1 |
| C5-G16 | N4-O6; N3-N1; O2-N2 |
| C5-A20 | O2-O2' |
| G6-C15 | N1-N3; N2-O2; O6-N4 |
| G6-A21 | O2'-OP1 |
| G7-C14 | N1-N3; N2-O2; O6-N4 |
| G7-A24 | N2-N1; N3-N6 |
| C8-G12 | N4-N7; N3-O6 |
| C8-A25 | O2-N6 |
| C8-C26 | O2-N4 |
| C10-G28 | N4-O6; N3-N1; O2-N2 |
| C11-G27 | N4-O6; N3-N1; O2-N2 |
| G12-C26 | N1-N3; N2-O2; O6-N4 |
| C14-A25 | O2'-N1; O2-N6 |
| C15-A23 | O2'-N1; O2-N6 |
| G16-A21 | N2-OP2; O2'-N7 |
| C17-A20 | O2-O2' |

We have determined the stability of the BWYV PK at 37 °C from its folding/unfolding equilibrium, using a structure-independent method described previously.^30^ Our simulation reproduces correctly a linear dependence of Δ*G* on ln *c* for *c* < 0.2 M, however it predicts an upward curvature of Δ*G* vs. ln *c* for *c* > 0.2 M (Fig. 2b). As is apparent below, this curvature can be traced to a sharp decrease in ΔΓ for large *c* which is due to the breakdown of the linearized Poisson-Boltzmann approximation. It is worth noting that even at *c* ≈ 0.5*M* the theoretical result for Δ*G* is only ≈ 7% higher the the experimental value. Because there is only one experimental data in the range *c* > 0.2 M (Fig. 2b), we cannot provide an accurate assessment of the magnitude of the discrepancies between simulation and experiment. There are two possibilities. (1) Given that the predicted heat capacity of the BWYV PK at 0.5 M is in near quantitative agreement with the experimental data (Fig. 2a) suggests that the measured Δ*G* at *c* ≈ 0.5*M* could have larger errors. (2) Alternatively, it is possible that small discrepancy between simulations and experiments does not imply the same behavior in the stability. Experiments for other RNAs and more data in the concentration range between 0.2*M* ≤ *c* ≤ 0.5 are needed to compare more precise comparisons between simulations and experiments.

**Figure 2:**
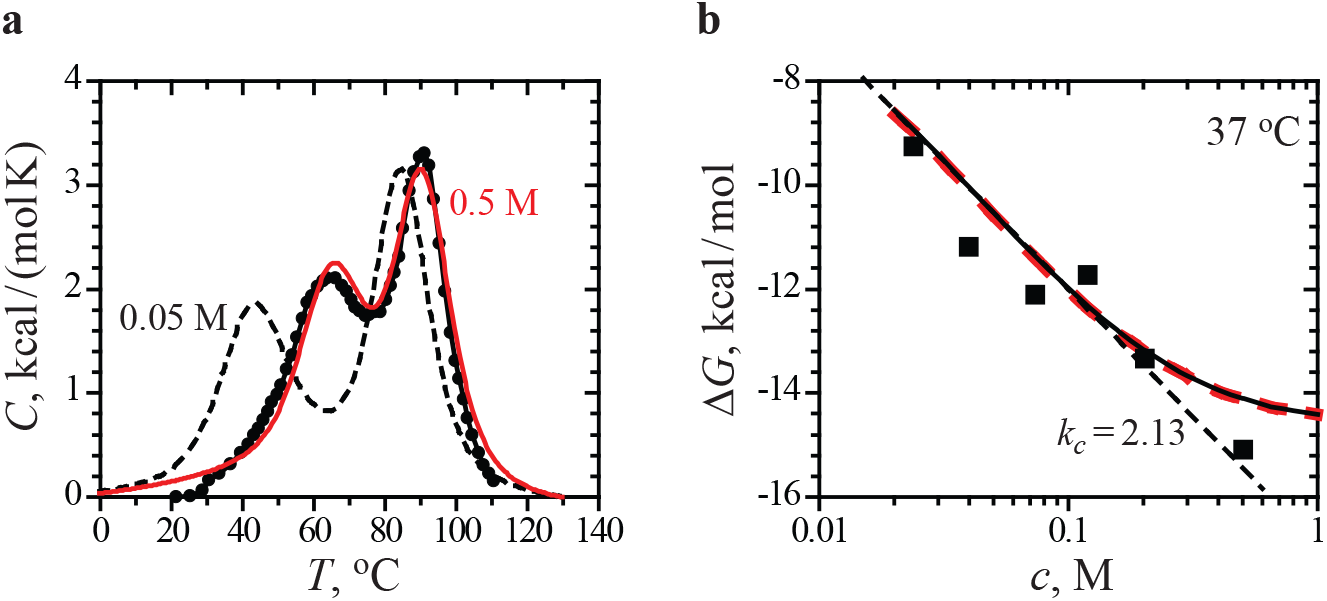
(a) Measured^20^ heat capacity, *C*, of the BWYV PK in 0.5 M K^+^ as a function of *T* (symbols). The solid curve (red) is the simulation data for *c* = 0.5 M. The dashed curve is the simulation data for *c* = 0.05 M (experimental data not available for comparison). The computed *C*(*T*) is plotted with respect to the heat capacity of the unfolded state at 130 °C. (b) Measured^14^ stability Δ*G* of the BWYV PK at 37 °C as a function of *c* (symbols). The solid curve is Δ*G* obtained from the analysis of the folding/unfolding equilibrium in simulations at various *c*. The thick dashed curve (red) is Δ*G*(*c*) = Δ*G*(c_0_) + ΔΔ*G*(*c*), where *c*_0_ = 0.2 M, Δ*G*(*c*_0_) is given by the solid curve, and ΔΔ*G*(*c*) is computed using Eq. (3) and ΔΓ(*c*) shown in Fig. 3b. *k_c_* specified in the figure panel is the slope of a linear fit of the simulation data for *c* < 0.2 M (straight dashed line).

**Figure 3:**
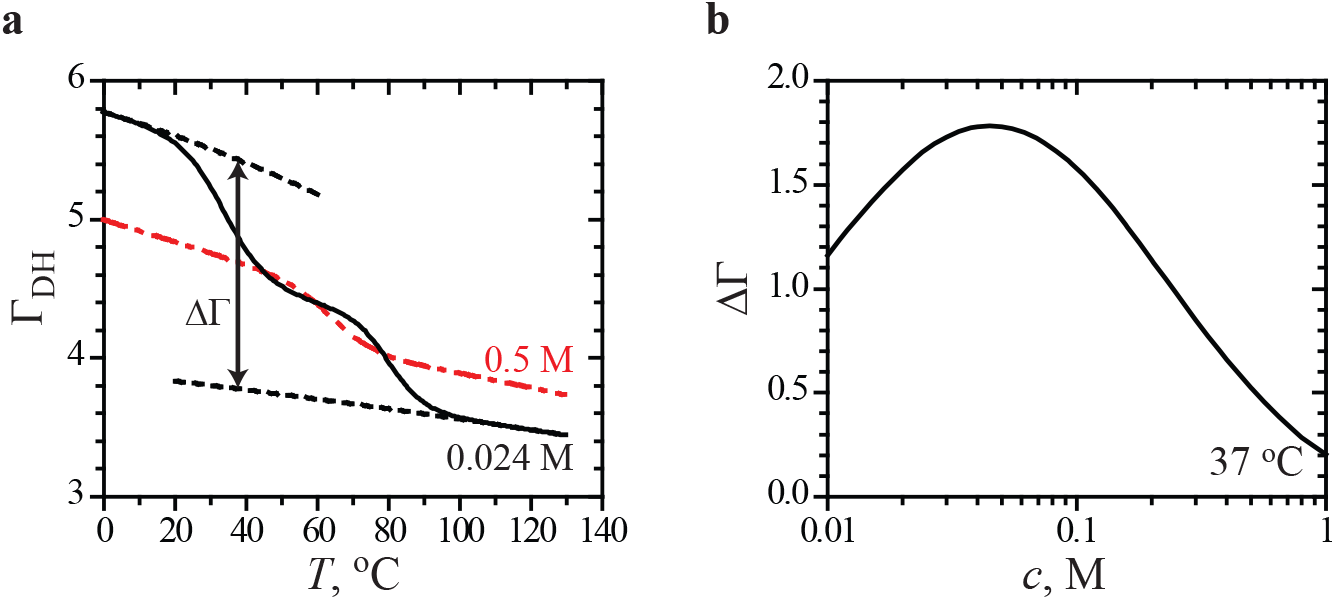
(a) Preferential interaction coefficient Γ_DH_, as a function of *T*, from simulations of the BWYV PK with *c* = 0.024 M (solid curve) and *c* = 0.5 M (dash-dotted curve, red). The dashed lines are the baselines representing the folded and unfolded states. The baselines are obtained by expanding Γ_DH_ to second order in *T* around *T* = 0 °C (folded) and *T* = 130 °C (unfolded). (b) ΔΓ = Γ_f_ – Γ_u_ at 37 °C as a function of *c*.

We now illustrate the connection between the preferential interaction coefficient Γ and stability of the BWYV PK. Figure 3a shows Γ_DH_ obtained from Eq. (9), where Γ_DH_ = Γ – (1 – *Q*)*N*_p_/2, as a function of *T*. The low and high *T* limits of this dependence are Γ_f_ – (1 – *Q*)*N*_p_/2 and Γ_u_ – (1 – *Q*)*N*_p_/2 in the fully folded and unfolded states, respectively. At intermediate tempera tures, ΔΓ = Γ_f_ – Γ_u_ is estimated as a difference between the “folded” and “unfolded” baselines plotted in Fig. 3a. We find that ΔΓ at 37 °C is a non-monotonic function of salt concentration *c* (Fig. 3b). Knowing Δ*G* at a reference *c*_0_ and ΔΓ(*c*), we can calculate Δ*G* at any *c* using Δ*G*(*c*) = Δ*G*(*c*_0_) + ΔΔ*G*(*c*) and Eq. (3). The result of this calculation at 37 °C is indistinguishable on the scale of the figure from Δ*G*(*c*) obtained directly from the folding/unfolding equilibrium (Fig. 2b). Interestingly, although ΔΓ(*c*) is a non-monotonic function of *c*, it yields an approximately linear dependence of Δ*G*(*c*) on ln *c* for *c* < 0.2 M. Therefore, linear fits of experimentally determined Δ*G*(*c*) do not give us an unambiguous estimate of ΔΓ. This point has already been discussed in the context of the nonlinear Poisson-Boltzmann equation for the BWYV PK in mixed Na^+^ and Mg^2+^ buffers, where the dependence of ΔΓ on the divalent ion concentration was also found to be non-monotonic.^14^

### Explicit ion simulations of the L10 HP

To our knowledge the simulations of the L10 HP reported here are the first computational study of RNA thermodynamics, besides our own studies of ribozyme folding^32^ using an explicit ion model. It was our intention to choose one of the most basic RNA molecules, such as the L10 HP, as a first test example. The same hairpin was also used as a benchmark for the development of an implicit ion model in our earlier work,^30^ because its thermodynamic stability was measured over a wide range of *c*.^22^ Because a high-resolution structure of the L10 HP is not available, we assume that the L10 HP native structure is an ideal A-form helix with six Watson-Crick base pairs and an unstructured loop.

We carried out grand-canonical simulations of the L10 HP in the temperature range from 0 to 130 °C for *c* = 0.05, 0.2 and 1 M. For every combination of *c* and *T*, the chemical potential of ions was determined from a canonical simulation of NaCl in the absence of RNA. The dependence of the chemical potential on *T* is found to be linear for all *c* (Fig. 4a), which permits a straightforward analysis of the grand-canonical data using weighted histograms.^30^ The L10 HP melting curves at three different *c* are characterized by a single melting peak, indicating cooperative melting of the hairpin stem (Fig. 4b). As anticipated, the position of the melting peak (or the melting temperature) increases substantially with ion concentration *c*.

**Figure 4:**
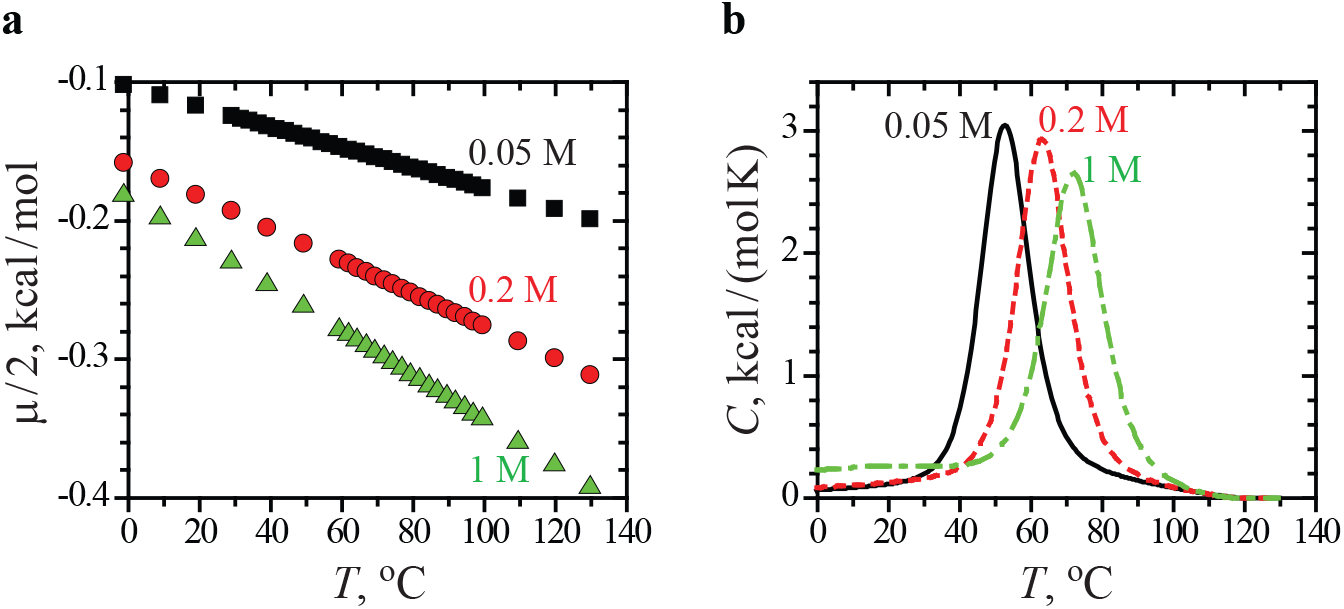
(a) Chemical potential of a neutral ion pair, *μ*, as a function of *T*, from explicit ion simulations of NaCl in the absence of RNA. Squares: *c* = 0.05 M, circles (red): *c* = 0.2 M, triangles (green): *c* =1 M. (b) Heat capacity, *C*, as a function of *T*, from grand-canonical simulations of the L10 HP in NaCl solution. The *C*(*T*) is plotted with respect to the heat capacity of the unfolded state. Solid line: *c* = 0.05 M, dashed (red): *c* = 0.2 M, dash-dotted (green): *c* =1 M.

We have determined the stability Δ*G*(*c*) of the L10 HP at 37 °C from its melting data using the structure-independent method described in our original paper.^30^ In the same paper we used this method to obtain Δ*G*(*c*) of the L10 HP at 37 °C from implicit ion simulations. The resulting Δ*G*(*c*) from the explicit and implicit ion models are compared in Fig. 5a.^30^ The explicit ion model yields a linear dependence of Δ*G* on ln *c* in the entire range of salt concentrations from 0.05 to 1 M. The linear fit of Δ*G* vs. ln *c* results in ΔΓ = 0.78 ± 0.07, which compares well with ΔΓ = 0.85 obtained by fitting the experimental data. However the actual values of Δ*G* are about 2 kcal/mol higher in the simulations than in experiment (Fig. 5a) at all *c*. The predictions of the implicit ion model compare more favorably with the experimental Δ*G*(*c*) in the range *c* < 0.2 M (Fig. 5a). This is because in the derivation of this model we used Δ*G*(*c*) of the L10 HP for calibration of the reduced RNA charge *Q*.^30^ For *c* > 0.2 M, the dependence of Δ*G* on ln *c* in the implicit ion model is strongly nonlinear due to the breakdown of the Debye-Hückel approximation, as in the case of the BWYV PK (Fig. 2b). In contrast, the linear behavior is captured by the model that models ion explicitly.

**Figure 5:**
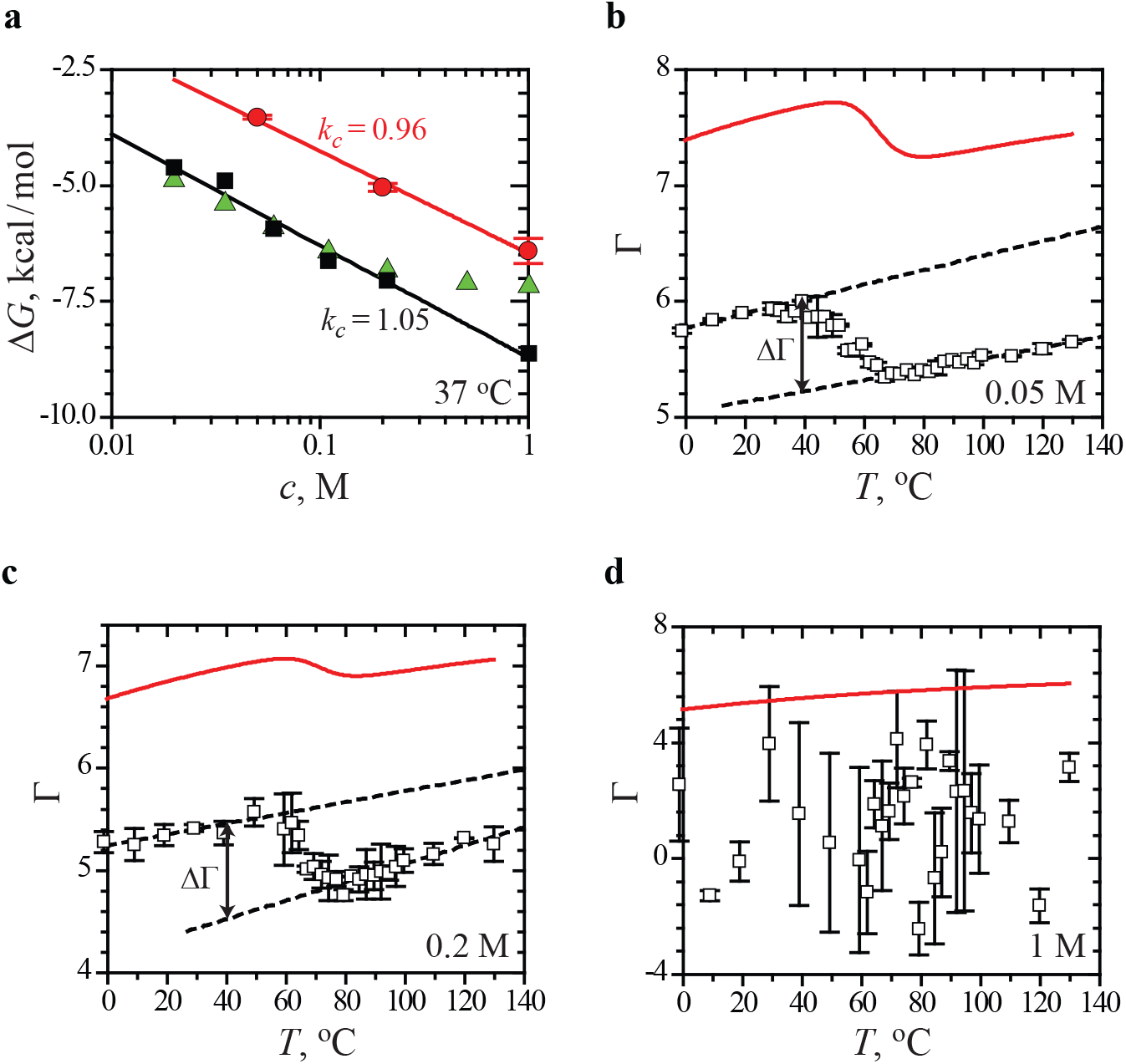
(a) Measured^22^ stability, Δ*G*, of the L10 HP at 37 °C as a function of *c* (squares). Circles (red) show Δ*G* obtained from the explicit ion simulations reported here. Triangles (green) show Δ*G* obtained from our implicit ion simulations reported previously.^30^ *k_c_* specified in the figure panel are the slopes of the linear fits of the experimental and explicit ion simulation data (solid lines). (b–d) Preferential interaction coefficient, Γ, as a function of *T*, from explicit (symbols) and implicit (solid curves, red) ion simulations of the L10 HP in NaCl solution. *c* is given in the figure panels. The dashed baselines in (b) and (c) are the least squares fits of the linear portions of the Γ vs. *T* curves. ΔΓ at 37 °C is defined as a difference between the low and high *T* baselines. Standard errors in Δ*G* and Γ were estimated by dividing the simulation data into two blocks and computing these quantities for each block.

The discrepancy in the estimates of Δ*G* obtained from the implicit and explicit ion simulations can be traced back to the preferential interaction coefficients Γ (Figs. 5b–d). For explicit ions, we define Γ = 0.5 (*N*_Na_ + *N*_Cl_) – *N*_NaCl_, where *N*_Na_ and *N*_Cl_ are the average numbers of Na^+^ and Cl^-^ ions in a grand-canonical simulation of an RNA molecule in NaCl solution with concentration *c* and volume *V*, and *N*_NaCl_ = 6.022 × 10^23^*cV*. The preferential interaction coefficient for implicit ions is defined by Eqs. (9) and (14). We find that the explicit ion model consistently underestimates Γ for all considered *c* and *T* (Figs. 5b–d). Because RNA molecules depend on counterion condensation for their ability to fold, the smaller values of Γ translate to higher free energies Δ*G*. For explicit ion simulations at *c* = 0.05 and 0.2 M, we were able to estimate ΔΓ at 37 °C from the dependence of Γ on *T* as illustrated in Figs. 5b and 5c. The resulting values, ΔΓ = 0.79 ± 0.08 at 0.05 M and 0.9 ± 0.4 at 0.2 M, agree within the error bars with ΔΓ extracted directly from the melting data. A large noise in Γ at 1 M prevented us from estimating ΔΓ for this salt concentration (Fig. 5d). Very long simulation times would be required to sufficiently reduce the noise level because typical ΔΓ are 3 orders of magnitude smaller than *N*_NaCl_ at 1 M (*N*_NaCl_ = 2032 for the simulation box with side 150 Å).

The stability data in Fig. 5a reveal the relative advantages and disadvantages of the implicit and explicit ion models. The implicit ion model essentially has one more free parameter, the reduced RNA charge *Q*. As demonstrated by our results presented here and previously,^30^ a single choice of *Q* works well for a variety of RNA molecules and solution conditions. Therefore, the implicit ion model may in fact be a preferred method if accurate estimates of Δ*G* are sought in the range *c* < 0.2 M. For larger salt concentrations, the dependence of Δ*G* on ln *c* develops what appears to be a false curvature and the model becomes less accurate.

By contrast, in the explicit ion model, the reduced RNA charge is fixed at –1 and is not a free parameter. The model consistently underestimates the value of Γ, the effect which appears to be independent of *T* and thus RNA conformation (Figs. 5b–d). We conjecture that this is a result of using a dielectric constant of water in Eq. (15), which is not expected to be very accurate in the vicinity of an RNA molecule that has a smaller dielectric constant. This can lead to an underestimation of the number of ions directly associated with the RNA backbone. The explicit ion model can still be useful in determining the form of the dependence of Δ*G* on ln *c*, if not the actual values of Δ*G*, since it does not break down at high salt concentrations *c* > 0.2 M. Let us also emphasize that neither BWYV PK nor L10 HP are stabilized by site-specific interactions with monovalent ions, and therefore both the implicit and explicit ion descriptions are applicable. In the case when site-specific interactions are present, for example in ribozymes, only the explicit ion model will provide correct structural information on the folded or partially folded states of the RNA.^32^

Direct comparison of the explicit and implicit ion simulation data in Fig. 5 shows that differences in Γ of 1.5–2 ions per RNA result in substantial differences in Δ*G*. Therefore, it becomes apparent that if any force field were to yield accurate estimates for Δ*G* it would have to, in the first place, provide highly accurate estimates for Γ. Here it is important to remember that Γ does not include only the ions directly bound to RNA, but it is defined as the total increase in the number of ions caused by an introduction of RNA in the simulation box. This and the long-range nature of the Coulomb potential effectively makes Γ a long-range property, whose accurate determination will require large simulation boxes to avoid finite-size effects. The necessity for large simulation boxes (or long simulation times) is just one technical difficulty that may affect an accurate estimation of RNA thermodynamics in simulations. A more general question is if it is possible to devise a universal force field, which includes only generic ion-RNA interactions and no structure-specific adjustable parameters, that would yield the correct ion numbers for diverse RNA systems. Our present results show that in the absence of such a model, simulations with implicit ions appears to be a reasonably good option currently available, where applicable, for the determination of RNA thermodynamics using simulations. We should also note that despite limitations (lack of ion-ion correlations and treating ions as point charges) apparently Γ values are well predicted using Non-Linear Poisson-Boltzmann (NLPB) equation. However, calculating Δ*G*(*c*) accurately requires sampling the ensemble of conformations of the folded, unfolded and intermediates (if any). Clearly, this requires simulations of the kind performed here.

## Discussion

### Effective Charge on the Phosphate Groups for RNA

Contrary to our expectations, we found that the implicit ion version of the model yields more accurate estimates for the RNA stability, Δ*G* as a function of ion concentration for L10 HP. A key parameter determining Δ*G* is the effective charge of phosphate groups, *Q*. In implicit ion simulations *Q* is calculated using the counterion condensation theory for an infinite cylinder with the mean axial distance between phosphate charges, *b*.^35^ By comparing with experimental data, we determined the optimal value of the free parameter *b* to be 4.4 Å.^30^ This result can be placed in the context of previous applications of counterion condensation theory to order-disorder transitions in polymeric nucleic acids.^34,36,37,39^

In these applications, all nucleic acid structures were treated as infinite cylinders, and the main distinction between the structures comprising a different number of strands was the mean distance, b, between the charges on the polymeric nucleic acids. Double-stranded and triple-stranded helices were characterized by 1.4 Å < *b* < 1.7 Å and *b* =1 Å, respectively.^34^ The formalism predicted 3.2 Å < *b* < 4.2 Å for a single-stranded (rod-like) nucleic acid polymer.^34,36,37^ By contrast, we used one value of *b* to describe both the folded and unfolded RNA in our simulations. Because distance 4.4 Å is consistent with previous estimates of *b* for single-stranded nucleic acids, we assume that it describes the geometry of the unfolded state of RNA reasonably well. This is further supported by our result that *b* = 4.4 Å works equally well for hairpins and pseudoknots that have significantly different charge densities in the folded state. We argue that specifying single *b* is sufficient in simulations, because the concept of counterion condensation must be invoked only in the unfolded state. Indeed, in simulations, RNA are flexible and the total charge density due to RNA and ions fluctuates in response to conformational changes. Although this version of the model does not include ions explicitly, they are effectively taken into account through the Debye-Hückel screening clouds in Eq. (7). As RNA folds, the backbone phosphates come in proximity, which results in a substantial overlap of the ionic clouds. Thus, any increase in phosphate charge density or Γ upon folding is taken into account through the conformational properties of the model itself. Good agreement with experiment indicates that the linear superposition of individual screening clouds is a valid model for an increased counterion uptake due to a conformational change. We speculate that the linear approximation, and thus the implicit ion model, will be quantitatively accurate for all folding transitions identified by small ΔΓ.

### Importance of RNA conformational ensemble and ion-ion correlation effects

Beyond the linear approximation, the electrostatic free energy of nucleic acids has been assessed by solving the nonlinear Poisson-Boltzmann equation using static fixed representations of the folded and unfolded states.^14,40–42^ In this case, the accuracy of the model is limited by the accuracy of representing the ensemble of unfolded states, which is typically described as a single conformation rather than a dynamic ensemble of conformations. When it comes to the study of RNA folding using simulations, it is more practical to use an explicit ion simulation model than to solve the NLPB equation at each time step. In the explicit ion model presented here, *Q* = –1 and there are no free parameters that may be varied to effectively tune ion-RNA interactions. We find that, if a single dielectric constant of water is used for all electrostatic interactions, the predictions of the explicit ion model for Δ*G* are quantitatively less accurate than those of the implicit ion model. Although it was not done here, one way to improve the explicit ion model would be to use a distance-dependent dielectric constant in Eq. (15). We emphasize that, despite their shortcomings, explicit ion simulations are the only valid way to study the effects of ion size, many body ion-ion correlations, or site specific ion-RNA interaction on nucleic acid properties, as we recently demonstrated for ribozymes, pseudoknots, and hairpins.^32^

### Uptake of ions as RNA folds and implications for single molecule pulling experiments

The logarithmic variation of Δ*G*(*c*) with monovalent ion concentration has previously been demonstrated using optical tweezer experiments^23^ only for RNA hairpins. Such experiments have been performed for RNA pseudoknots at specific salt concentrations. It would be most interesting to perform them over a range of concentrations in order to most directly measure Δ*G*(*c*) as a function of *c* in order to test the predictions made here for the BWYV pseudoknot.

It is remarkable that the difference between ion uptake per RNA between the folded and unfolded states is only between 1.5–2 (Fig. 5), which is is not that dissimilar to other RNA molecules.^43^ The small difference in ΔΓ implies that there are a large number of ions that are territorially bound to RNA even in the unfolded state, thus affecting both the enthalpy and entropy of the disordered RNA. Although ensemble experiments have been used to obtain accurate estimates of it appears that single molecule pulling experiments^44^ combined with simulations^45^ are efficacious in directly obtaining the *c*-dependence of ΔΓ. The experimental tests of our predictions for the differential uptake of ions between the folded and unfolded states are most readily done using single molecule pulling experiments.^44,46,47^

## Conclusions

We have presented and tested a coarse-grained model to study the folding thermodynamics of RNA in implicit and explicit ion simulations. Our model provides the first unified framework to obtain the RNA thermodynamic stability and ion preferential interaction coefficient entirely from simulations, without resorting to any experimental measurements. Here, we performed an independent validation of the analytical relationship between these two quantities. Because the model and the protocols are general they be used to calculate accurately the folding thermodynamics of arbitrary RNA, thereby providing the much needed insights into how ions control their self-assembly.

## Methods

### RNA model with implicit ionic buffer

All details of the coarse-grained simulation model that do not pertain to electrostatic interactions can be found in our earlier paper.^30^ We assume that the coarse-grained sites representing RNA bases and sugars have no charge. The charge on each coarse-grained phosphate group is taken to be –*Qe*, where *e* is the proton charge and *Q* is smaller than 1 due to partial charge neutralization by cations due to ion condensation. We determine the value of the reduced charge on the phosphate groups using Manning’s theory of counterion condensation,^35^ as was done previously for the *Tetrahymena* ribozyme.^3^ For an infinitely long rod-like polyelectrolyte, with charge –*e* per contour length *b*_0_, Manning’s theory predicts^35^

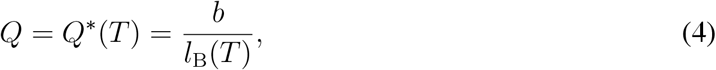

where *b* = *b*_0_/*z*_c_, *z*_c_ is the valence of screening cations, and *l*_B_ is the Bjerrum length,

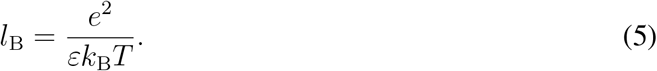

The reduced charge *Q* depends on the temperature *T* nonlinearly because the dielectric constant of water, *ε*, decreases with *T* as^48^

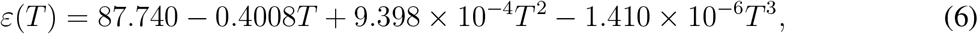

where *T* is in units of °C. We have previously shown that simulations using *b* = *b*_0_ = 4.4 Å in Eq. (4) reproduce the measured stabilities and melting profiles of different RNA molecules, including the L10 HP, over a wide range of monovalent salt concentration.^30^ Our results below demonstrate that *b* = 4.4 Å provides good agreement with experimental data for the BWYV PK as well.

The uncondensed ions, which presumably exchange with condensed ions, in the simulation model are described by the Debye-Hückel, theory. For a given conformation of an RNA molecule in monovalent salt solution, the electrostatic free energy in the Debye-Hückel approximation, *G*_DH_, is^40^

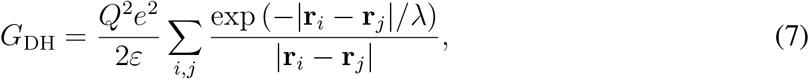

where **r**_*i*_, **r**_*j*_ are the coordinates of phosphates *i* and *j*, λ is the Debye-Hückel screening length,

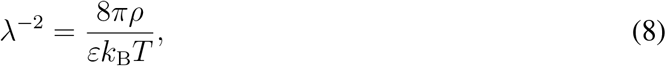

and *ρ* is the bulk number density of counterions or coions, *ρ* = 6.022 × 10^23^*c*. The corresponding expression for the preferential interaction coefficient in the Debye-Hückel approximation, Γ_DH_, is

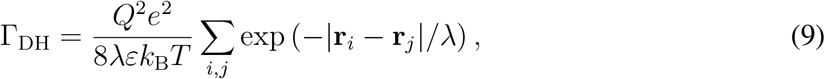

where **r**_*i*_, **r**_*j*_ are the same as in Eq. (7). One way to derive Eq. (9) is from the spatially varying number densities of counterions and coions,

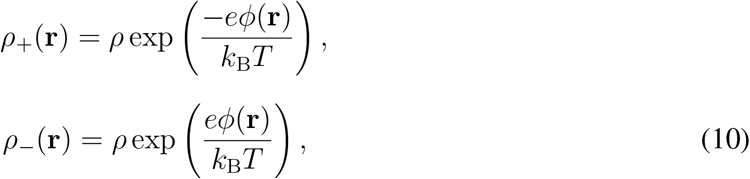

where *ϕ*(**r**) is the total electrostatic potential at **r** due to the ions and phosphates *i*,

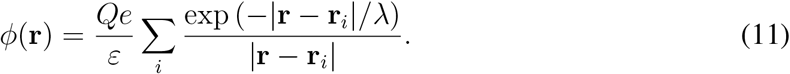

By expressing Γ_DH_ as the space integral,

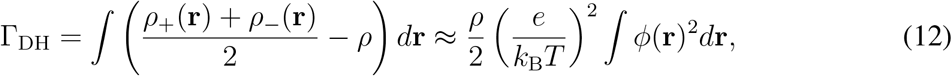

and substituting Eq. (11) in Eq. (12), we recover Eq. (9). Equation (9) can also be obtained from

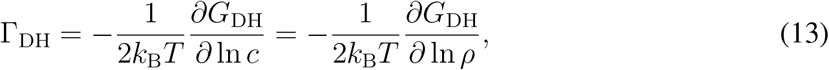

where *G*_DH_ is given by Eq. (7).

In simulations of the BWYV PK we use *G*_DH_ and *∂G*_DH_/*∂***r**_*i*_ to compute the electrostatic energy and forces between phosphate groups, thus implicitly taking into account the ionic buffer. The preferential interaction coefficient is computed by averaging the expression in Eq. (9) over all sampled conformations at given *c* and *T*. We note that the full expression for the preferential interaction coefficient should also include the contribution from the condensed ions,

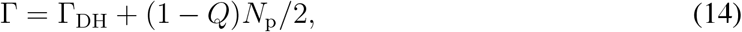

where *N*_p_ is the total number of phosphate groups in the RNA molecule. Because the condensed ion term is independent of RNA conformation it does not contribute to ΔΓ or the salt dependence of Δ*G*. However, it must be taken into account when comparing the preferential interaction coefficients obtained in implicit and explicit ion simulations.

### RNA model with explicit ionic buffer

In simulations of the L10 HP in NaCl solution, the Na^+^ and Cl^−^ ions are modeled explicitly as spheres with an appropriate charge, *Q_i_*, and radius, *R_i_*. Solvent molecules are not explicitly included in simulation. For two sites *i* and *j* with charges *Q_i_* and *Q_j_* (ions or phosphates), the electrostatic interaction as a function of distance *r* is modeled using the Coulomb potential *U_C_*,

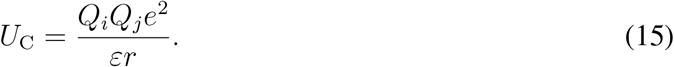

Ewald sums are used to compute the total electrostatic potential and force for each site. Because Manning’s theory should not be invoked when treating ions explicitly, the charge on phosphate groups *Q* = –1.

Excluded volume between any sites *i* and *j* is described by the modified Lennard-Jones potential,

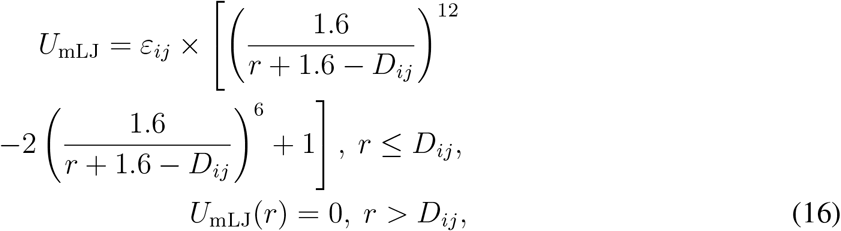

where *D_ij_* = *R_i_* + *R_j_* and 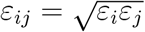.

The potentials *U*_C_, *U*_mLJ_ and parameters *Q_i_* were adopted from our earlier work,^32^ where we showed that RNA thermodynamics is relatively insensitive to the parametrization of monovalent ions. The ionic radius and *ε_i_* for sodium ion, which were not listed in the supplementary information in^32^ are *R_i_*=1.868Å, *ε_i_* = 0.00277*kcalmol*^-1^, respectively. All the remaining elements of the force field are carried over from the implicit ion model.^30^

The explicit ion simulations are implemented in the grand ensemble for Na^+^ and Cl^−^ with the purpose of determining the preferential interaction coefficient Γ. For given *c* and *T*, we first compute the chemical potential of a neutral ion pair in a canonical simulation of NaCl in the absence of RNA using the Widom insertion technique. Once the chemical potential is known, it is used in a grand-canonical simulation of the L10 HP in NaCl. In the grand-canonical simulation a single attempt to add or remove an ion pair is made at each time step, and the preferential interaction coefficient is obtained by simply averaging the excess number of ions in the simulation box.

Finally, as discussed in the context of our original model,^30^ it is only possible to use a weighted histogram technique for data analysis if the interaction potentials are either independent or linearly dependent on *T*. The Coulomb potential *U*_C_ in Eq. (15) depends on *T* nonlinearly through the dielectric constant *ε*. Therefore, to be able to use weighted histograms, the *U*_C_ actually employed in simulations was expanded to first order in *T* around *T* = 50 °C (in the middle of the relevant temperature range). The same linear expansion is also applied to *Q* in Eq. (14) for direct comparison of Γ obtained from the implicit and explicit ion models.

### General simulation details

The dynamics of RNA and ions were simulated by solving the Langevin equation of motion. In explicit ion simulations a single L10 HP molecule was contained in a cubic box with side 150 Å, and the number of Na^+^ and Cl^−^ ions was calculated using *c* and the simulation box volume. A single trajectory was generated for all considered *c* and *T* in implicit and explicit ion simulations. The simulation trajectories were started in the folded state and were at least several *μ*s long. This length was sufficient to attain equilibrium at all conditions. Other details of Langevin dynamics simulations, which are the same for implicit and explicit ion models, are given in our earlier paper.^30^ The reader is referred to the same paper for a description of data analysis techniques, including a structure-independent method to calculate the RNA stability Δ*G*.

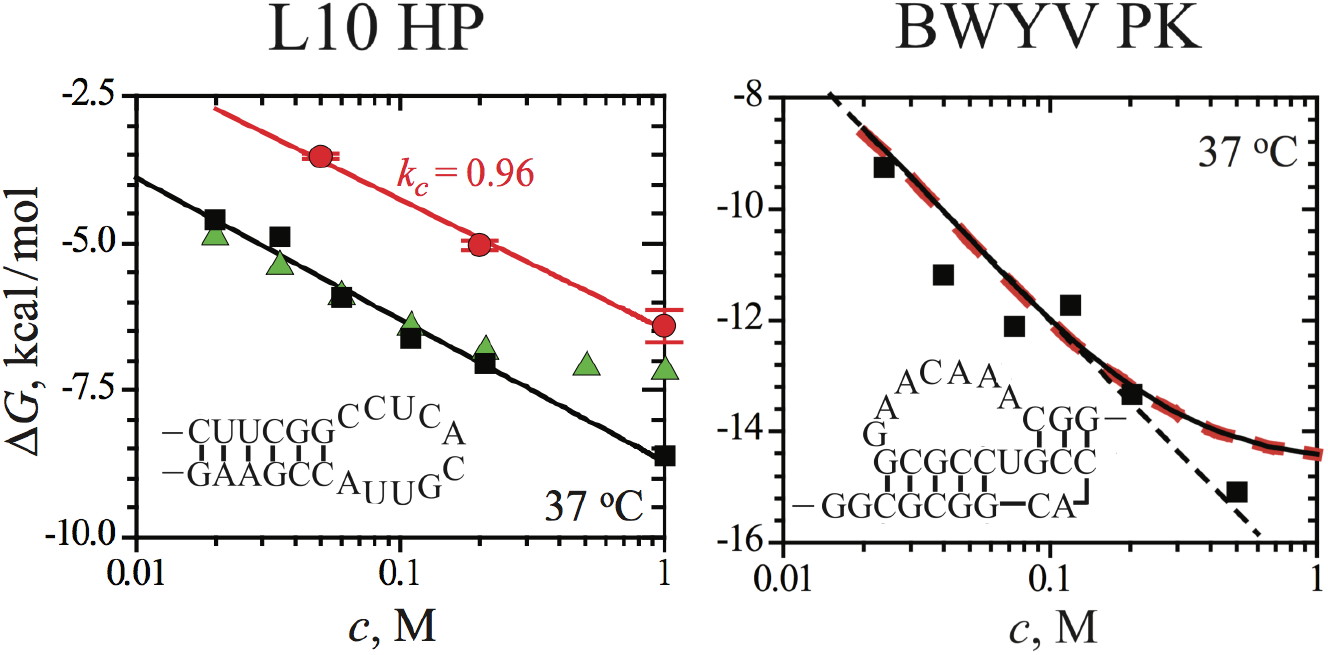

## Acknowledgements

We thank Naoto Hori for useful discussions and help with graphics. This work, which was completed when the authors were in the Institute for Physical Science and Technology, was supported by a grant from the National Science Foundation (CHE 16-36424) and the Welch Foundation (F-0019) through the Collie-Welch Chair.

